# PV-specific loss of the transcriptional coactivator PGC-1α slows down the evolution of epileptic activity in an acute ictogenic model

**DOI:** 10.1101/2021.06.25.449915

**Authors:** R. Ryley Parrish, Connie Mackenzie-Gray-Scott, Darren Walsh, Claudia Racca, Rita M. Cowell, Andrew J. Trevelyan

## Abstract

The transcriptional coactivator, PGC-1α (peroxisome proliferator activated receptor gamma coactivator 1α), plays a key role coordinating energy requirement within cells. Its importance is reflected in the growing number of psychiatric and neurological conditions that have been associated with reduced PGC-1α levels. In cortical networks, PGC-1α is required for the induction of parvalbumin (PV) expression in interneurons, and PGC-1α deficiency affects synchronous GABAergic release. It is unknown, however, how this affects cortical excitability. We show here that knocking down PGC-1α specifically in the PV-expressing cells (PGC-1α^PV-/-^), blocks the activity-dependent regulation of the synaptic proteins, *SYT2* and *CPLX1*. More surprisingly, this cell-class specific knock-out of PGC-1α appears to have a novel anti-epileptic effect, as assayed in brain slices bathed in 0 Mg^2+^ media. The rate of pre-ictal discharges developed approximately equivalently in wild-type and PGC-1α^PV-/-^ brain slices, but the intensity of these discharges was lower in PGC-1α^PV-/-^ slices, as evident from the reduced power in the gamma range and reduced firing rates in both PV interneurons and pyramidal cells during these discharges. Reflecting this reduced intensity in the pre-ictal discharges, the PGC-1α^PV-/-^ brain slices experienced many more discharges before transitioning into a seizure-like event. Consequently, there was a large increase in the latency to the first seizure-like event in brain slices lacking PGC-1α in PV interneurons. We conclude that knocking down PGC-1α limits the range of PV interneuron firing, and this slows the pathophysiological escalation during ictogenesis.

## Introduction

Studies in non-neuronal tissue have identified the transcriptional coactivator, PGC-1α (peroxisome proliferator activated receptor gamma coactivator 1α), as a key regulator of expression of nuclear encoded mitochondrial genes (Austin and St-Pierre, 2012; Handschin and Spiegelman, 2006; Lin et al., 2005; McMeekin et al., 2021). Its expression is induced by various triggers, including exercise, cold and fasting, and it then drives mitochondrial (Puigserver et al., 2001) and peroxisome biogenesis (Bagattin et al., 2010) and uncouples mitochondrial respiration (St-Pierre et al., 2003). PGC-1α is also highly expressed in different parts of the brain, where its expression is enriched in GABAergic neurons (Cowell et al., 2007). Within cortical networks, PGC-1α appears particularly to affect parvalbumin-expressing (PV) interneurons (Cowell et al., 2007): the highest transcriptomic counts of PGC-1α are found in this subset of cortical neurons (Saunders et al., 2018); reduced PGC-1α expression (heterozygotes and homozygote knock-out) leads to a loss of PV in a dose-dependent manner, without affecting the expression of other interneuronal markers, while overexpression of PGC-1α in cultured neurons strongly induces PV expression (Lucas et al., 2010). Consistent with these findings, PV interneurons ordinarily have a very high mitochondrial content, and express cytochrome c more intensely than other cortical neurons (Gulyas et al., 2006; Kageyama and Wong-Riley, 1982). PV interneurons are also among the most active of any cortical neuronal class; they characteristically fire on every cycle of gamma oscillations (Atallah and Scanziani, 2009; Hajos et al., 2004; Whittington and Traub, 2003) and show extremely high firing rates ahead of an ictal wavefront (Parrish et al., 2019; Sessolo et al., 2015). These various considerations – their apparent high metabolism, and the powerful effect that PV interneurons exert upon cortical networks – have led to the suggestion that PV interneurons may be particularly susceptible to metabolic stress, which in turn could give rise to neuropathology (Kann, 2016; Whittaker et al., 2011). It is relevant, therefore, that reduced PGC-1α expression has been found in a number of neurological conditions (McMeekin et al., 2021), including Parkinson’s disease (Zheng et al., 2010), Alzheimer’s disease (Qin et al., 2009; Sheng et al., 2012), Huntington’s disease (Cui et al., 2006), multiple sclerosis (Witte et al., 2013), and schizophrenia (Christoforou et al., 2007; Jiang et al., 2013a), but interestingly, not epilepsy. Germline knockdown of PGC-1α is associated with hyperactive behaviour (Lin et al., 2004), although this is believed to be a behavioural adaptation to a reduction in thermogenesis capacity in brown fat and muscle, because the hyperactive phenotype was not replicated in mice with central nervous system deletion of PGC-1α (Lucas et al., 2012).

Knocking-out PGC-1α selectively in PV interneurons (PGC-1α^PV-/-^) also affects synaptic function, with a marked drop in two synaptic proteins, synaptotagmin 2 (Syt2) and complexin 1 (Cplx1), a reduction in synchronous GABA release, and impaired long-term memory (Lucas et al., 2014). Intriguingly, the effects on GABAergic synaptic function were somewhat inconsistent in hippocampal and neocortical networks (Dougherty et al., 2014; Lucas et al., 2014; Lucas et al., 2010). These changes might be expected to alter network excitability, so we investigated how epileptiform activity develops in brain slices prepared from these mice, compared to age-matched wild-type controls, when bathed in artificial cerebro-spinal fluid (aCSF) lacking Mg^2+^ ions (0Mg^2+^ model). Both experimental groups showed preictal activity occurring at similar rates, but the intensity of these discharges, as measured by the gamma bandwidth power, and the mean firing rates in both PV and pyramidal neurons, was lower in the brain slices prepared from PGC-1α^PV-/-^ mice. Perhaps reflecting this lower intensity of discharge, the knock-out brain slices experienced many more preictal events prior to the first seizure-like event (SLE), and consequently, also showed a significantly longer latency until this first SLE. Altogether, these data indicate that lack of PGC-1α within inhibitory PV cells acts as a restraint on runaway excitation within cortical networks, having a neuroprotective effect during induced epileptogenesis. This result demonstrates a role for PGC-1α in influencing cortical excitability through transcriptional regulation of PV cell activity.

## Methods

### Animals

All procedures were performed according to the guidelines of the Home Office UK and animals (Scientific Procedures) Act 1986. All the data were collected from adult male and female mice (2 – 6 months). Wild type (C57/BL6) mice or PVCreHet mice (Stock # 008069, The Jackson Laboratory, Bar Harbor, ME USA) were used as control animals for all experiments. No difference was observed in the 0 Mg^2+^ induced activity between wild type and PVCreHet mice, so these groups were combined and labelled “WT” in the study. Conditional deletion of PGC-1α within PV interneurons was achieved by crossing mice with LoxP sites flanking the exon 3–5 region of the PGC-1α gene (PGC-1α ^fl/fl^, Stock # 009666, The Jackson Laboratory) with mice expressing Cre recombinase driven by PV (mentioned above) on a C57BL/6 background on which they were maintained. Briefly, a line of homozygous PGC-1α^fl/fl^ mice were created and crossed with a homozygous PVCre mouse line to generate the experimental animals with the genotype PVCre:PGC-1α^fl/+^. All animals were group housed in individually ventilated cages kept at room temperature with a 12hour/12hour light/dark cycle and provided with food and water *ad libitum*.

### Viral injections

To allow visualization of PV interneurons for the patch clamp experiments, 2-4 month old PVCre or PGC-1α.PVCre animals were injected with AAV9.Floxxed.MCherry viral vector (Penn Vector Core, University of Pennsylvania). Mice were anaesthetised by intraperitoneal injection of ketamine (75 mg kg^-1^, Ketalar™ Injection, Pfizer Ltd., Sandwich, UK) combined with medetomidine (1 mg kg^-1^, Domitor®, Janssen Animal Health, Basingstoke, UK) and were then transferred to a stereotaxic frame where anaesthesia was maintained using inhalational administration of isofluorane (3-5% for induction and 2% for maintenance, delivered in 02 at 400-800ml/min). The head was shaved and disinfected with chlorhexidine. A craniotomy was performed to reveal the dorsal right cortex, and two injections were made of the viral vector (1 in 10 dilution) separated along the antero-posterior axis (~1mm and 2mm posterior to bregma) and at the same mediolateral location (~1.5-2mm lateral to the midline). The viral vector was delivered by pressure injection (Ultramicropump 3, World Precision Instruments), through a 10μl Hamilton syringe, with a bevelled 36 gauge needle (World Precision Instruments). At each site, a total of 1μl was injected at a rate of 5-10nl/s, at 3 depths (200 / 400 / 400nl at 1 / 0.7 / 0.4mm, respectively, deep to the pia). The needle was left *in situ* for 3 mins after the final injection, to prevent back-tracking of the injected particles out of the injection track. Post-operatively, mice were given subcutaneous injection of atipamezole (5 mg kg^−1^, Janssen Animal Health) and meloxicam (5 mg kg^-1^) to reverse the anaesthesia and provide post-operative pain relief. The animals were allowed to recover in a heated and ventilated incubation chamber overnight and monitored at 2 and 24 hours post-surgery. Some animals were given further doses of meloxicam and fluids, as required, based upon clinical assessment of their post-operative recovery, using body weight and behavioural scoring. Animals were subsequently assessed for their general health twice a day, for 5 days, and then weekly for 2-5 more weeks, prior to being sacrificed to prepare brain slices.

### Brain slice preparation

Mice were sacrificed by cervical dislocation followed by immediate decapitation. The brain was removed and sliced into ice-cold cutting solution containing (in mM): (3 MgCl_2_; 126 NaCl; 2.6 NaHCO_3_; 3.5 KCl; 1.26 NaH_2_PO_4_; 10 glucose), using a Leica VT1200 vibratome (Nussloch, Germany). We used 400μm horizontal sections, containing entorhinal cortex (EC) and neocortex, for local field potential (LFP) recordings, and 350μm coronal sections for the patch clamp recordings. Slices were then transferred to an interface tissue holding chamber and incubated for 1 - 2 hours at room temperature in aCSF containing (in mM): (2 CaCl_2_; 1 MgCl_2_; 126 NaCl; 2.6 NaHCO_3_; 3.5 KCl; 1.26 NaH2PO4; 10 glucose). All the solutions were being bubbled continuously to saturate with carboxygen (95% O_2_ and 5% CO_2_).

### Extracellular LFP recordings

These experiments were performed using interface recording chambers. Slices were placed in the recording chamber perfused with aCSF or aCSF without Mg^2+^. Recordings were obtained using normal aCSF-filled 1-3MΩ borosilicate glass microelectrodes (GC120TF-10; Harvard apparatus, Kent) placed in deep layers of the neocortex. Microelectrodes were pulled using Narishige electrode puller (Narishige Scientific Instruments, Tokyo, Japan). The temperature of the chamber and perfusate was maintained at 33-36 °C using a closed circulating heater (FH16D, Grant instruments, Cambridge, UK). The solutions were perfused at the rate of 3-4mls/min by a peristaltic pump Watson Marlow 501U (Watson-Marlow Pumps Limited, Cornwall UK). Waveform signals were acquired using BMA-931 biopotential amplifier (Dataq instruments, Akron, USA), Micro 1401-3 ADC board (Cambridge Electronic Design, UK) and Spike2 version 7.10 software (Cambridge Electronic Design, UK). Signals were sampled at 10 kHz, amplified (gain: 500) and bandpass filtered (1-3000Hz). CED4001-16 Mains Pulser (Cambridge Electronic Design, UK) was connected to the events input of CED micro 1401-3 ADC board and was used to remove 50Hz hum offline. Extracellular field recordings were analysed to detect pathological discharges using a custom-written code in Matlab2018b (Mathworks, MA, USA) which can be provided upon request. In brief, this used a frequency-domain analysis to detect periods when the LFP power exceeded baseline activity by a designated threshold. Using the spectrogram.m macro, in Matlab (bin width 0.128s, with 50% time shifts), we derived the mean and standard deviation for an epoch of baseline lasting at least 10s, for frequencies between 1-150Hz (50 subdivisions, on a log scale). The spectrogram was then performed for the entire recording, and normalized to the baseline mean and standard deviation for each frequency bandwidth individually. To avoid contamination with mains noise, we omitted frequencies between 48-52Hz. This process generated a z-score spectrogram, in which the pixels represented the excess power at a particular frequency, for a specific time bin (0.128s), in terms of standard deviations from the baseline mean. Using a threshold of 3 standard deviations above the baseline mean, we identified all time bins for which any frequency exceeded this threshold. These were summed cumulatively, to estimate the rate and acceleration of pathological activity within the brain slice. Continuous epochs of suprathreshold bins were designated to be single “events”, which were then analysed for their total power within the conventional LFP frequency bins (α, γ bands etc) over the whole event.

### Patch-clamp recordings

These recordings were performed in submerged recording chambers. Slices were perfused with aCSF with the same composition as mentioned above at 3-5 mL/min and heated to 33–34°C. Whole cell and cell attached data were acquired using pClamp software, Multiclamp 700B, and Digidata acquisition board (Molecular Devices, Sunnyvale, CA, USA). Recording chamber, heater plate (Warner Instruments, Hamden, CT) and micromanipulators (Scientifica,, UK) were mounted on a Scientifica movable top plate fitted to a laser spinning disc-confocal microscope (Visitech, UK). Whole cell and cell attached recordings of layer five neurons within neocortex were made using 4–7 MΩ pipettes (GC150F-10, Harvard apparatus, Kent) pulled using micropipette puller (Model-P87, Sutter Instruments, CA, USA). Pipettes were filled with solution containing (mM): 125 K-gluconate, 6 NaCl, 10 HEPES, 2.5 Mg-ATP, 0.3 Na2-GTP; pH and osmolarity of the electrode-filling solution used was adjusted to 7.4 and 280-285 mOsms, respectively.

To investigate the firing properties of pyramidal cells and PV interneurons during induced epileptogenesis dual recordings were made in neocortical layer 5 cells in voltage clamp configuration. First a pyramidal cell was patched in whole cell mode and held at −70mV to act as a reference for the development of ictal-like events in the slice and secondly either a pyramidal or PV cell was patched in cell attached mode to identify spikes during evolving epileptiform activity as monitored by the whole cell reference recording. Once the whole cell and cell attached patch recordings were established, the Mg^2+^ ions were washed out of the bathing aCSF to induce the development of epileptiform activity. For the targeted cell attached patch recordings of PV interneurons the cells were visualised using a spinning disc confocal microscope (UMPlanFL N ×20, 0.5 NA objective) with a rhodamine (U-MRFPHQ) filter set. The tissue was illuminated with a 561 nm laser (Cobolt Jive 50; Cobolt) to visualize the MCherry fluorescence within the PV interneurons of the injected PVCre and PGC-1α.PVCre animals. Hamamatsu C9100 EM cameras (Hamamatsu Photonics (UK), Welwyn Garden City, UK) were used to visualize and capture images, run by Simple PCI software (Digital Pixel, Brighton, UK; microscope 1) installed on a Dell Precision computer (Dell Technologies, Round Rock, TX, USA). Patch clamp data were analysed using custom-written codes in Matlab2018b.

### Tissue collection

After 1 hour of recordings using the interface chamber, tissue was immersed in Ambion RNAlater solution (Thermo Fisher Scientific) and the neocortical tissue was collected. Tissue was then stored in RNAlater solution, at 4 °C until the RNA was extracted

### RNA Preparations

RNA was extracted using the RNAqueous-Micro Total RNA Isolation Kit (Thermo Fisher Scientific) and eluted in 15 μl of provided elution buffer. Residual genomic DNA was removed using reagents supplied in the above kit and done according to the Manufacturer’s instructions. Concentration and purity of RNA were determined using a BioDrop DUO (BioDrop, Cambridge, UK). RNA was then stored at −80 ^°^C until the cDNA library preparation.

### Real time (RT) PCR

Samples were normalized to 100 ng prior to the cDNA library preparation. cDNA synthesis was preformed using the Applied Biosystems High-Capacity cDNA Reverse Transcription Kit (Thermo Fisher Scientific) in a Thermo Hybaid PCR Express Thermal Cycler (Thermo Hybaid, Ashford, UK) at a total volume of 20 μl. The cDNA was then diluted 1:4 with nuclease free H2O. The following primers were used for the RT-PCR: Parvalbumin (For: attgaggaggatgagctggg, Rev: cgagaaggactgagatgggg), c-fos (For: cagcctttcctactaccattcc, Rev: acagatctgcgcaaaagtcc), Glyceraldehyde 3-phosphate dehydrogenase (Gapdh) (For: acctttgatgctggggctggc, Rev: gggctgagttgggatggggact). Ppargc1α (For: agcctctttgcccagatctt, Rev: ggcaatccgtcttcatccac), Syt2 (For: caactcccattcctgaccct, Rev: ttttctccaccgcccatttg), Cplx1 (For: tcagacacacttggagtccc, Rev: cacttgattatgcggccctc). Q-PCR amplifications were performed on a Applied Biosystems Step One Plus RT PCR system (Thermo Fisher Scientific) at 95 °C for 10min, 40 repeats of 95 °C for 10s, followed by 58 °C for 30s, 95 °C for 15s. This protocol was immediately followed by the melt curve starting at 95 °C for 15s, 60 °C for 1min, 95 °C for 15s, and, finally, held at 4 °C. All samples were run in duplicate at a primer molarity of 10 μmol/L, and PV, c-fos, Ppargc1α, Syt2, Cplx1 were compared to GAPDH. Expression of GAPDH was unchanged across all treatment groups. Cycle threshold (Ct) values were analysed using the comparative Ct method to calculate differences in gene expression between samples. Samples were run on a DNA agarose gel to confirm expected product size.

### Statistical analysis

Relative mRNA fold changes were analysed using the comparative Ct method. Due to a lack of homogeneity between groups (Brown–Forsythe test and Bartlett’s test), mRNA expression data were first transformed using a square root transformation. Square root transformation of mRNA data resulted in a nonsignificant Brown–Forsythe test and Bartlett’s test. Transformed data were then analysed using a one-way analysis of variance (ANOVA), with a Tukey post-hoc test for all mRNA expression data of 3 or more groups. Electrophysiology event detection data and mRNA of only two groups were analysed using the student’s t-test. Significance was set at P ≤ 0.05 for all analyses and data represented as mean ± SEM unless stated otherwise. Figures of mRNA expression and event detection, along with all statistics were done using GraphPad Prism (GraphPad Software, Inc., La Jolla, California, USA). Figures of electrophysiology traces were created using Matlab R2018b (MathWorks, USA), Inkscape and Adobe Photoshop.

## Results

Seizure activity rapidly induces a strong expression of proto-oncogenes, including c-fos. This has been demonstrated in a range of *in vivo* models of ictogenesis (Dragunow and Robertson, 1987; Morgan et al., 1987; White and Gall, 1987). Similarly, in brain slices, following an hour of exposure to 0 Mg^2+^ aCSF, c-fos and PGC-1α are strongly up-regulated, as are the synaptic proteins, synaptotagmin 2 (*Syt2*) and complexin 1 (*Cplx1*), while *Pvalb* is significantly down-regulated (Figure 1A). The relevance of these changes in *Syt2, Cplx1* and *Pvalb* is that expression of all three have previously been shown to be influenced by PGC-1α levels (Lucas et al., 2014; Lucas et al., 2010). In PGC-1α^PV-/-^ brain slices, *Pvalb, Syt2* and *Cplx1* all show greatly reduced expression in the naïve slices (pre-exposure to 0Mg^2+^ aCSF) compared to wild-type slices, and critically, showed no alteration of expression following exposure to 0 Mg^2+^ aCSF (Figure 1B-D), indicating that the activity-induced changes in these genes in wild-type slices are indeed downstream of PGC-1α.

**Figure 1.**
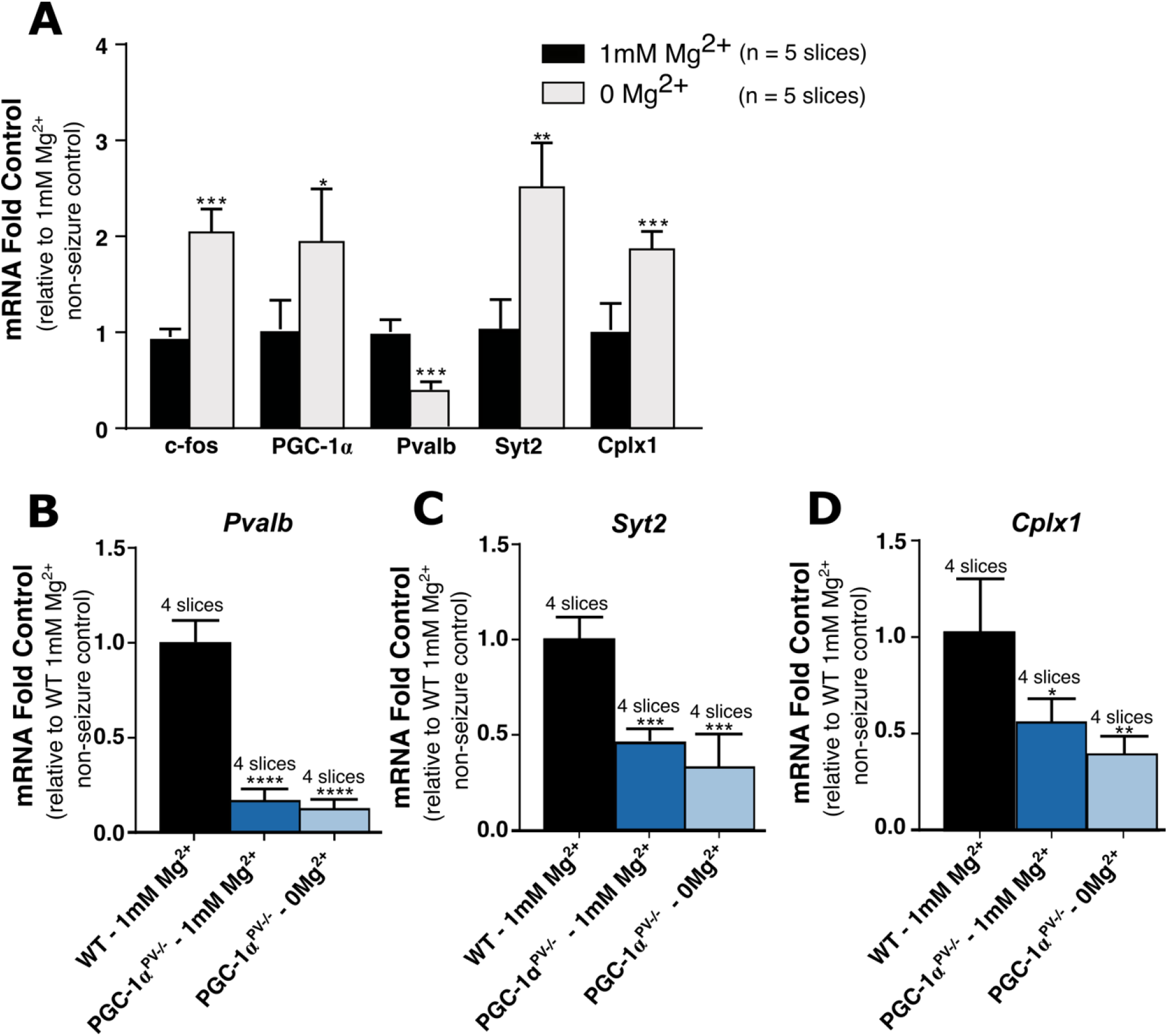
Ictogenesis induced gene-expression changes are markedly altered in PGC-1α^PV-/-^ mice. (A) Shows the increased expression of c-fos, PGC-1α, parvalbumin (Pvalb), synaptotagmin 2 (Syt2) and complexin 1 (Cplx1) after 1 hour being bathed in 0mM Mg^2+^ ACSF, relative to non-seizure control (1mM Mg^2+^). (**B-D**) mRNA levels for parvalbumin (*Pvalb*) **(B**), synaptotagmin 2 (*Syt2*) **(C**) and complexin 1 (*Cplx1*) (**D**), normalised to the mean for wild-type brain slices kept 1hr in normal 1mM Mg^2+^ ACSF (“control”, black). The relative change in mRNA is shown also for brain slices, prepared from mice with PGC-1α knocked-out in PV cells, and bathed either in normal 1mM Mg^2+^ (dark blue) or in 0mM Mg^2+^ ACSF (light blue). * p < 0.05; ** p < 0.01; *** p < 0.001; **** p <0.0001.

The lack of change in gene expression in PGC-1α^PV-/-^ brain slices was not due to a lack of epileptiform activity during the hour exposure to 0Mg^2+^ aCSF; indeed, both sets of brain slices showed the distinctive pattern of evolving pathological discharges (Figure 2A), starting with transient discharges lasting a few hundred milliseconds, and at longer latency, the appearance of sustained rhythmic discharges lasting tens of seconds, which we term seizure-like events (SLEs). The cumulative activity patterns, however, were strikingly different for the two experimental groups (Figure 2B-E). PGC-1α^PV-/-^ brain slices experienced a significantly larger number of preictal events prior to the first SLE (Figure 2B) relative to control brain slices, which cumulatively, represented a much longer duration of pathological activity (Figure 2C). The first SLE happened significantly later in PGC-1α^PV-/-^ brain slices (Figure 2D), while the total number of SLEs within the first 30mins exposure to 0Mg^2+^ aCSF was significantly less, compared to wild-type slices (Figure 2E). The cumulative duration of pathological activity rose steadily in both experimental groups, but because there were so many more preictal events in PGC-1α^PV-/-^ brain slices, the cumulative total in that experimental group achieved a far higher level prior to the first SLE (Figure 3A). Notably, however, the discharge durations remained stable in both groups (Figure 3B; note the constant gradients), and furthermore, there was no difference in gradient between the groups.

**Figure 2.**
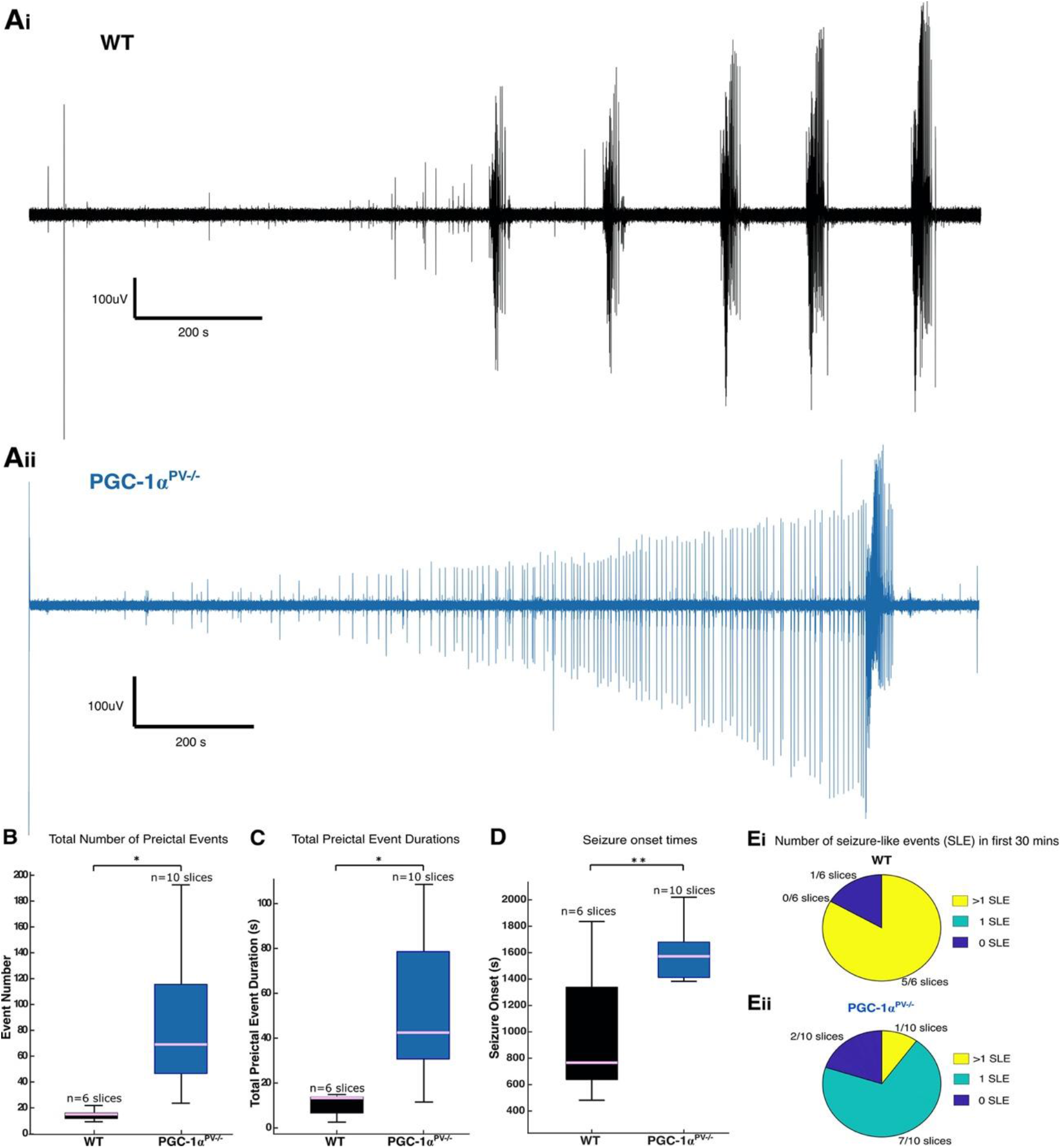
Markedly different patterns of epileptiform activity leading up to the first SLE, in brain slices prepared from PGC-1α and wild-type mice. (A) Representative LFP recordings of neocortical activity during 0Mg^2+^-induced ictogenesis in brain slices prepared either from wild-type (Ai) or PGC-1α^PV-/-^ mice (Aii). (B) Boxplot showing the total number of preictal events (events prior to first seizure-like event (SLE; pink line = median; boxed region = interquartile range; whiskers – full data range) for wild-type (black) and PGC-1α^PV-/-^ mice (blue). (C) Boxplot showing the total cumulative duration of pathological preictal activity. (D) Boxplot showing the latency to the first SLE. (E) Pie charts showing the number of SLEs in the first 30 minutes in 0Mg^2+^ for wild-type (Ei) and PGC-1α^PV-/-^ mice (Eii). No SLEs (dark blue), one SLE (turquoise) and more than one SLE (yellow). * p < 0.05; ** p < 0.01.

**Figure 3.**
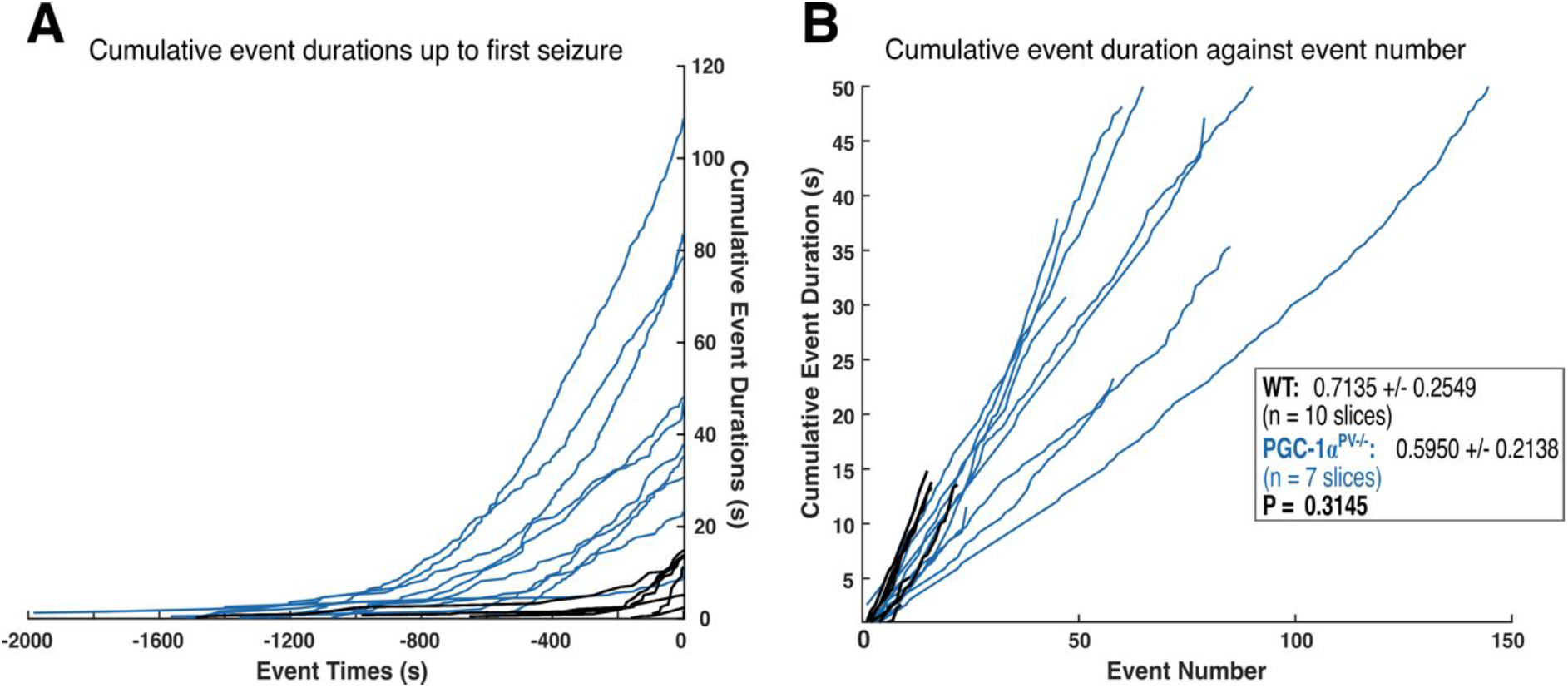
PGC-1α^PV-/-^ mice show far greater cumulative pathological activity prior to the onset of SLEs, compared to wild-type mice. (A) The cumulative total duration of the preictal pathological activity leading up to the onset of the first seizure-like event (SLE, designated t = 0s). (B) Shows the cumulative events durations with progressing event number for wild-type (black) and PGC-1α^PV-/-^ mice (blue). Note the steady gradients (inset box), indicative that the event durations do not change appreciably over time.

We next analysed the power spectra of each preictal discharge (Figure 4), which showed that although the durations remain stable, the power across all bandwidths increased progressively, leading up to the first SLE. The low frequency component of the LFP is thought to reflect the local synaptic currents, while the high gamma activity scales with the level of local firing. The ratio of these two bandwidths, therefore, is a measure of the degree to which synaptic volleys induce local postsynaptic firing. The gamma / delta ratio for the wild-type brain slices was significantly larger than recorded in the PGC-1α^PV-/-^ brain slices, when all pre-ictal events were included (Figure 4C,D), and the difference between groups was maintained when the analysis was restricted to the events in the 100s prior to the SLE (Figure 4E,F) (there was a fractional drop in the gamma/delta ratio for the PGC-1α^PV-/-^ brain slices in this final 100s).

**Figure 4.**
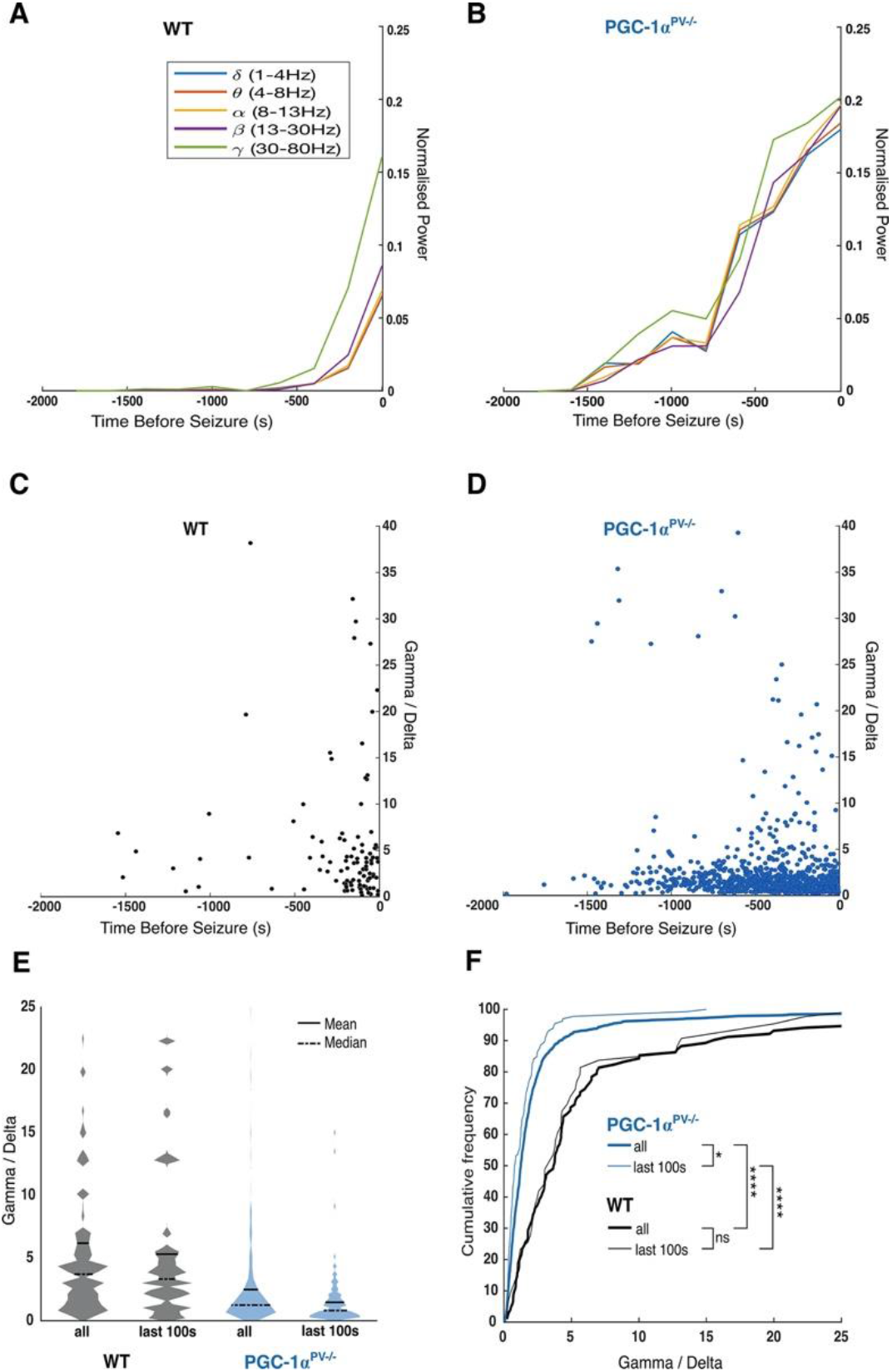
Preictal discharges in brain slices from PGC-1α knock-out mice show reduced normalised gamma activity (γ/δ ratio), compared to wild-type mice. (**A**) The mean normalised power, for the main physiological LFP bandwidths as indicated, averaged for sequential 200s epochs prior to the onset of the first SLE (t = 0s), for wild-type brain slices, recorded in 0 Mg^2+^ ACSF (n = 7). (**B**) Equivalent data from brain slices from PGC-1α^PV-/-^ mice (n = 10). Note the far more rapid escalation of activity in the wild-type brain slices, especially in the gamma and high-gamma frequency bands, leading up to the first SLE. (**C**) The gamma/delta ratio for every single interictal event, plotted against time relative to the onset of the first SLE, in the wild-type brain slices, and (**D**) in the PGC-1α^PV-/-^ brain slices. (**E**) Violin plots of γ/δ ratios, for the entire data set of interictal events, and also events occurring in the 100s immediately prior to the first SLE. (**F**) Cumulative frequency plots of the same data sets. Note the highly significant difference between WT and knock-out tissue, and the fact that there is minimal, if any, change in the period immediately prior to the SLE. * p < 0.05; **** p < 0.0001.

To investigate these differences further, we made paired patch clamp recordings from PV interneurons and pyramidal cells (Figure 5). The pyramidal recording was made in whole cell voltage-clamp mode, to record the synaptic bombardment, while the PV recording was made in cell-attached mode, to record the firing pattern (Figure 5Ai,Bi). This allowed us to compare the level of PV participation in these discharges, in the wild-type and PGC-1α^PV-/-^ experimental groups. We found that PV firing rates were significantly lower in the PGC-1α^PV-/-^ brain slices compared with wild-type brain slices prior to the first SLE (Figure 6Ai), although this difference was not maintained in later interictal events (Figure 6Aii). The number of action potentials per discharge (Figure 6B), and the action potential durations (Figure 6C), were equivalent in the wild-type and PGC-1α^PV-/-^ brain slices.

**Figure 5.**
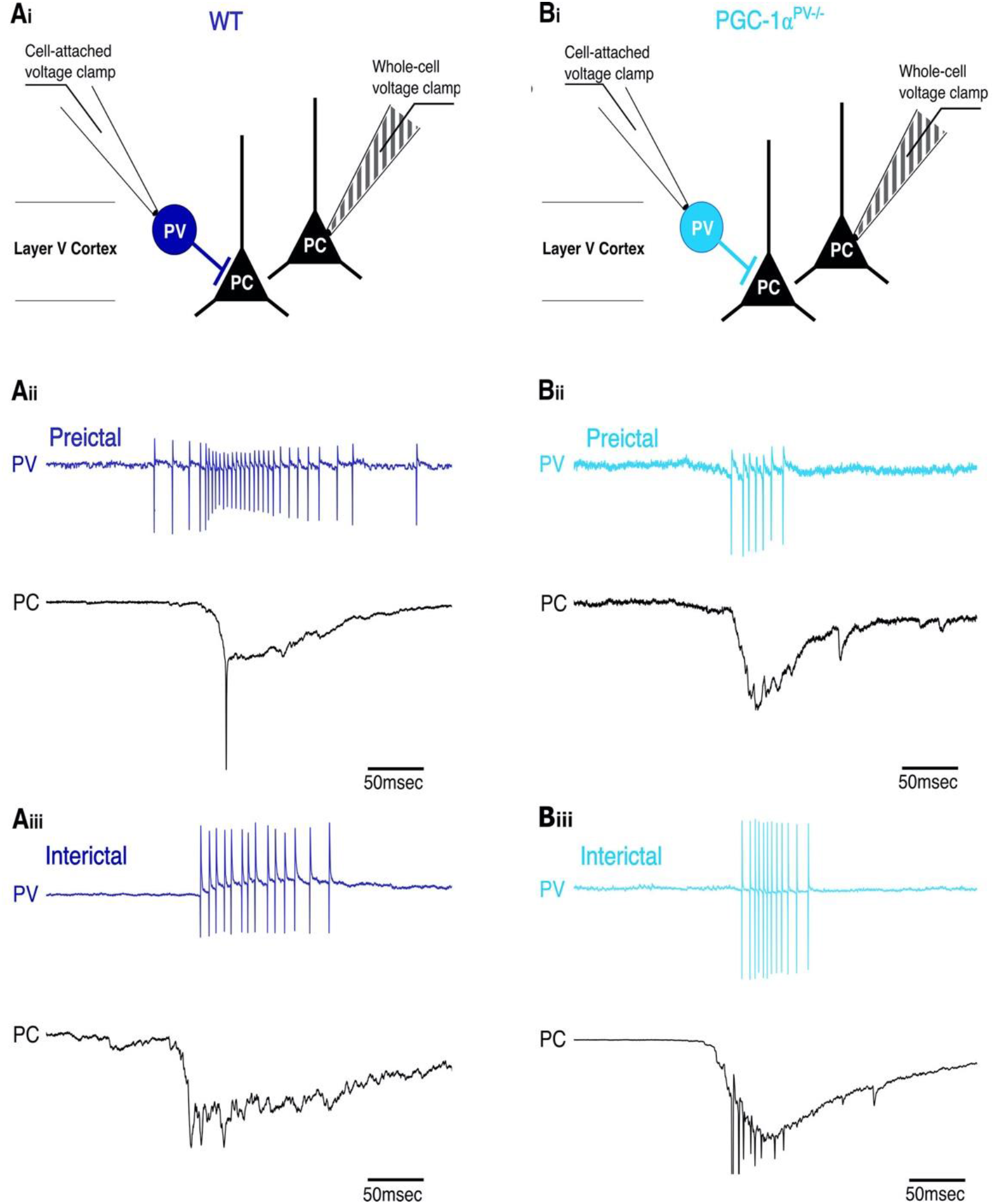
Example PV interneuron preictal and interictal events. (**Ai** and **Bi**) The recording configuration for the example traces, below, with a whole-cell, voltage-clamp, pyramidal cell (triangle) recording (shown by the striped electrode) and a cell-attached, voltage-clamp PV interneuron (circle) recording (shown by the clear electrode). The dark red indicates WT and the orange indicates PGC-1α^PV-/-^ mice. The underlying panels show preictal (**Aii**, **Bii**), and interictal (**Aiii**, **Biii**) pyramidal cell and interneuron events. The top traces show the cell-attached recording and the lower traces show the whole-cell recording. WT group, dark blue; PGC-1α^PV-/-^ group, light blue.

**Figure 6.**
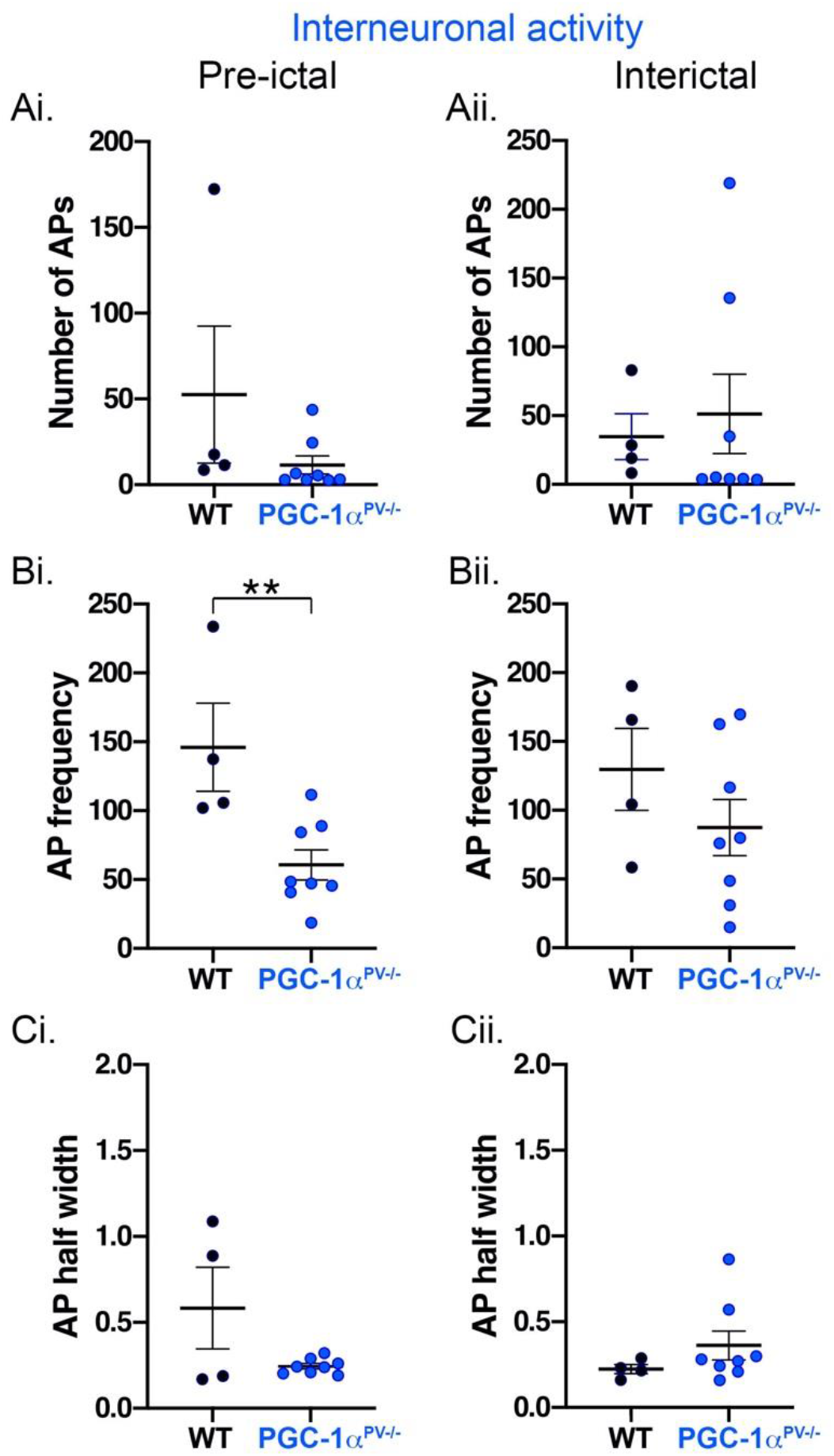
Action potential properties of PV interneurons during preictal and interictal events following 0Mg^2+^-induced ictogenesis. This figure shows cortical PV interneuron action potential numbers (**Ai**, **Aii**), their frequency (**Bi**, **Bii**) and halfwidths (**Ci**, **Cii**) during preictal (top row) and interictal events (bottom row) induced by 0Mg^2+^-ictogenic aCSF in control (black) and PGC-1α^PV-/-^ (blue) mouse tissue. Data points show the average data per brain slice. ** p < 0.01.

We then repeated the experiment, but instead, recording the firing patterns of the pyramidal cells (Figure 7). The numbers of action potentials per discharge was similar for the two experimental groups (Figure 8A), but the maximal firing rate for the PGC-1α^PV-/-^ brain slices was significantly lower than for the wild-type brain slices, consistent with the lower gamma/delta ratio (Figure 8B). This difference was seen when analyses included all interictal pathological discharges (Figure 8Bii), or if the analysis was restricted just to the events prior to the first SLE (Figure 8Bi). There was no difference in the half-width of the action potentials in the two groups (Figure 8C). These analyses show, therefore, that even though the genetic manipulation was restricted to the PV interneuronal cell class, the network impact was felt more widely across the network.

**Figure 7.**
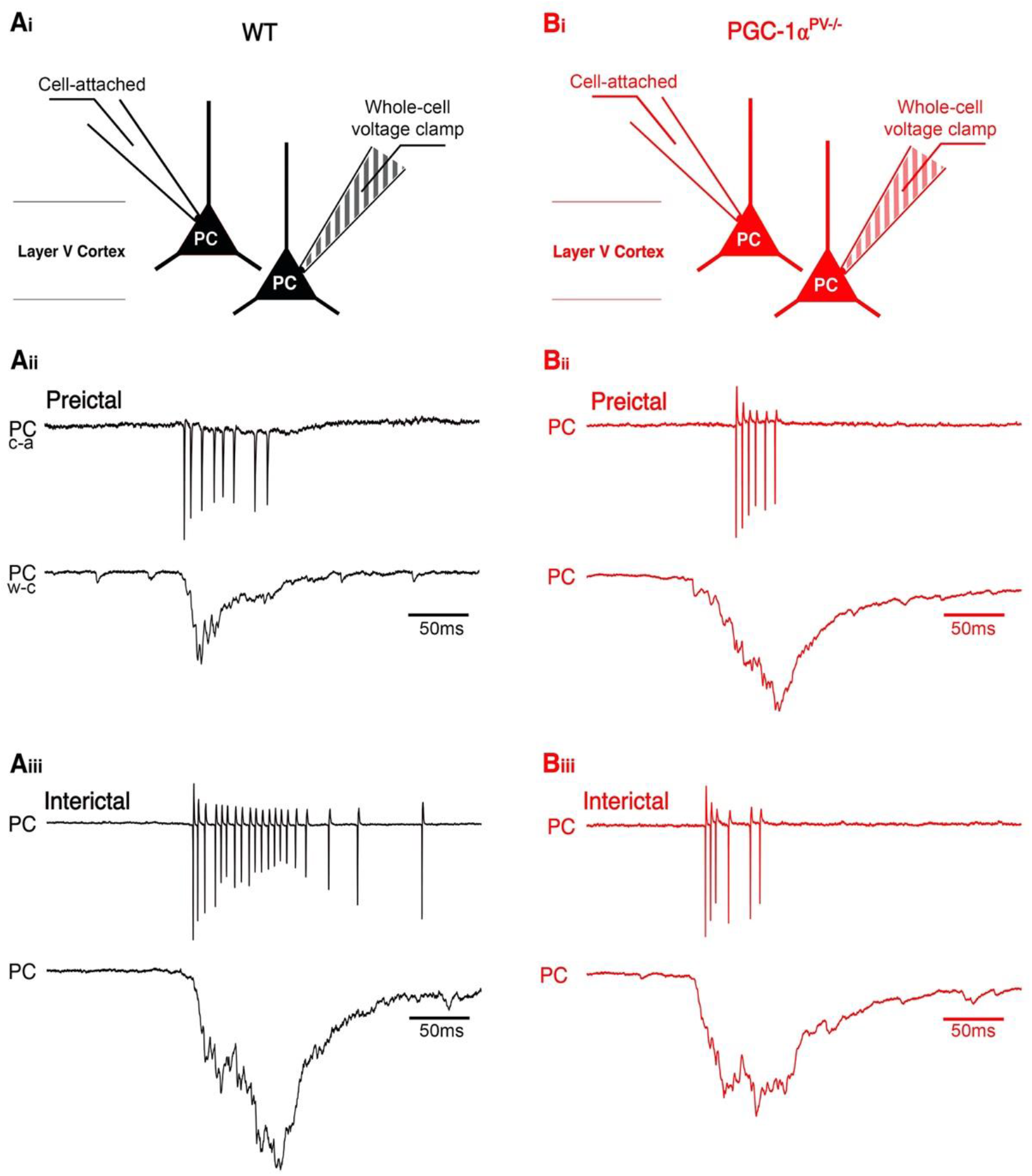
PGC-1α^PV-/-^ mice show reduced pyramidal firing rates, during preictal discharges, compared with wild-type mice. (Ai and Bi) The recording configuration for the underlying traces. The dark blue indicates WT and the light blue indicates PGC-1α^PV-/-^ pyramidal cells. The underlying panels show preictal (Aii, Bii), and interictal (Aiii, Biii) pyramidal cell events. The top traces show the cell-attached recording and the lower traces show the whole-cell recording.

**Figure 8.**
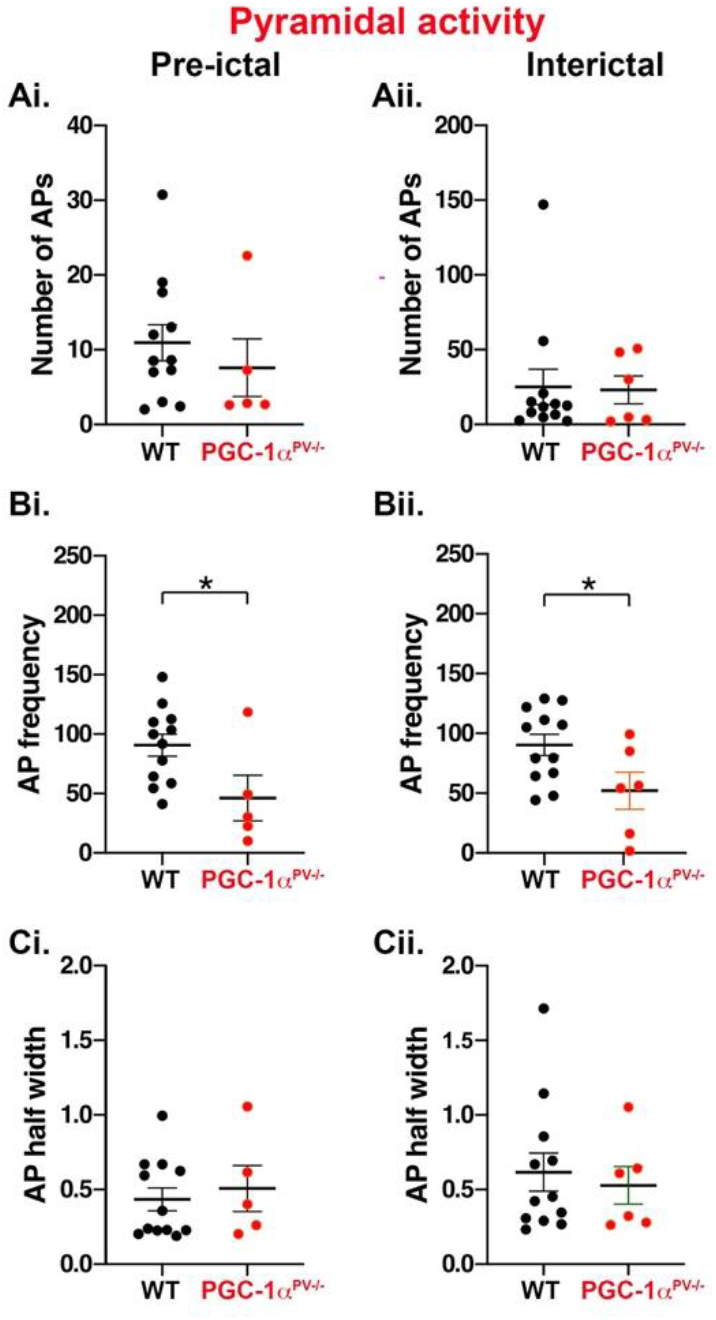
Action potential properties of pyramidal cells during preictal and interictal events following 0Mg^2+^-induced ictogenesis. This figure shows cortical pyramidal cell action potential numbers (**Ai**, **Aii**), their frequency (**Bi**, **Bii**) and halfwidths (**Ci**, **Cii**) during preictal (top row) and interictal events (bottom row) induced by 0Mg^2+^-ictogenic aCSF in control (black) and PGC-1α^PV-/-^ (red) mouse tissue. Data points show the average data per brain slice. * p < 0.05.

## Discussion

We have shown in these studies that knocking down the transcriptional coactivator, PGC-1α, specifically in PV interneurons, reduces network excitability. Our focus upon this subpopulation of interneurons was motivated in part because they have been postulated to play pivotal roles in various physiological brain rhythms (Cobb et al., 1995; Sohal et al., 2009; Traub and Miles, 1991), as well as both opposing (Parrish et al., 2019; Trevelyan, 2016), and promoting (Chang et al., 2018), seizure-like activity in different situations (note that these two statements are not necessarily incompatible – see (Burman et al., 2019; Cossart et al., 2005; Ellender et al., 2014; Sessolo et al., 2015)). Finally, there is a close relationship between PGC-1α and PV expression (Lucas et al., 2010), for which we provide further evidence, by showing that activity dependent regulation of genes downstream of PGC-1α such as PV is lost in the knock out (Figure 1 B, C and D). The anti-epileptic effect reflects a reduction in the intensity of pre-ictal bursts, measured both by the high-gamma power, and also by the firing rates of both PV interneurons and pyramidal cells.

The effect on PV interneuronal firing could be understood as a direct consequence of the fact that these are the cells in which PGC-1α is knocked-down. As such, it could be interpreted as an adaptation to a mutation which limits energy production, by limiting also the energy expenditure. However, the secondary, effect on pyramidal firing tells us that there are indirect consequences of the cell-specific knock-down, and given that interneuronal firing is driven by pyramidal activity, there are clearly more complex feedback loops that cloud any simple interpretation of the results. This is further complicated by other effects, such as the reduction in expression of genes downstream of PGC-1α; synaptotagmin 2 (Syt2) and complexin 1 (Cplx1). Syt2 and Cplx1 are synaptic proteins which act to promote synchronous neurotransmitter release from PV cells and their expression levels are reduced in PGC-1α^PV-/-^ mice (Lucas et al., 2014). We confirmed this reduction and additionally showed a lack of activity-dependent regulation of expression levels which resulted in a failure to increase expression in response to induced hyperexcitability.

Another interesting element is the reduction also in PV expression in PGC-1α^PV-/-^ mice. PV acts as a slow calcium buffer, helping to protect against hyperexcitability by sequestering away excess Ca^2+^ during periods of intense activity. Altered Ca^2+^ buffering in PGC-1α^PV-/-^ mice may therefore be a factor in the reduced level of interneuronal bursting, although it is notable that the specific knock-down of PV in striatal interneurons is associated with increased excitability (Orduz et al., 2013), the opposite of what we found during induced ictogenesis. The different results may arise for several reasons: our studies involved a reduction in PV expression, rather than a complete PV deficiency; they were of neocortical, rather than striatal interneurons; and the activity patterns and input to PV cells during 0Mg^2+^ induced ictogenesis may differ substantially from those of electrical stimulation paradigms, and as already mentioned, are influenced by multiple other factors besides.

The nonautonomous cellular effect seen in pyramidal cells may also arise through a complex set of interactions. One possibility is suggested by the very rapid escalation of interictal activity intensity in the wild-type experimental group, once it starts (see Figure 4A), relative to that seen in the PGC-1α^PV-/-^ brain slices (Figure 4B). The rate of escalation in epileptiform activity can be explained in terms of the rate of chloride loading, which in turn, depends on the rate of PV firing. Lowering the PV activity, as is seen in the PGC-1α^PV-/-^ brain slices, would reduce this effect, slowing the escalation, and when averaged over multiple preictal events, yields a lower mean discharge rate also for the pyramidal population. In contrast, the intense GABAergic discharges seen in wild-type brain slices, while serving a protective purpose also has the unwanted consequence of loading their postsynaptic target cells with chloride (Thompson and Gahwiler, 1989), thereby contributing to the pathological evolution of activity (Alfonsa et al., 2015; Burman et al., 2019; Cossart et al., 2005; Dzhala et al., 2010; Ellender et al., 2014; Huberfeld et al., 2007; Lillis et al., 2012). This is an example of positive feedback, where raised network activity creates the substrate that facilitates further neuronal activation, that may be critical for seizure initiation (Graham et al., 2021).

Early studies of PGC-1α identified its thermogenic role in skeletal muscle and brown fat, by uncoupling mitochondrial respiration in response to cold (Puigserver et al., 1998); interestingly, cold does not induce PGC-1α in the brain. There, its role may relate to protection against other stresses, including cytokines and lipopolysaccharides (McMeekin et al., 2021; Puigserver et al., 2001). For instance, PGC-1α leads to increases in expression of various enzymes involved in detoxification of reactive oxygen species (Austin and St-Pierre, 2012). Conversely, in mouse models of stress, PGC-1α reduction associated with increased markers of oxidative stress, are found within cortical PV interneurons (Jiang et al., 2013b). The cellular mechanism appears to involve activation of the p38 MAPK pathway, leading to phosphorylation of PGC-1α, which stabilizes the protein and increases its transcription inducing activity (Puigserver et al., 2001). It is noteworthy therefore, that inhibition of the p38 MAPK pathway has also been reported to increase the latency to the first seizure, following pilocarpine treatment (Zhou et al., 2020). That is to say, the pharmacological blockade of MAPK and the knockdown of one of its targets, PGC-1α, as we show in this study, appear to have similar effects, both slowing the pathological process in acute models of ictogenesis. Inhibiting p38 MAPK also reduces the frequency of seizures, and the level of associated pathological damage within the hippocampus (Zhou et al., 2020). In contrast, periods of status epilepticus activate p38 MAPK, and this may mediate the subsequent anatomical changes within the cortical networks (Hu et al., 2020).

Reduced PGC-1α has been associated with a range of neurological conditions (Christoforou et al., 2007; Cui et al., 2006; Jiang et al., 2013a; Qin et al., 2009; Sheng et al., 2012; Witte et al., 2013; Zheng et al., 2010), but not, to date, with epilepsy. The most pertinent of these to our study is the association with schizophrenia (Christoforou et al., 2007; Jiang et al., 2013a), for which the pathophysiological role of PGC-1α is also believed to be manifest through the PV interneurons within cortical networks (Bartley et al., 2015; Jiang et al., 2013a). The clinical association between epilepsy and schizophrenia is well established, with the existence of one condition carrying around a 6-8-fold increase in risk for having the other (Chang et al., 2011). Previous work reported that knocking down PGC-1α in PV interneurons alters the asynchronous synaptic release of GABA (Lucas et al., 2014), which may be expected to alter how PV interneurons influence spike timing and information flow through cortical networks. Our work suggests that different forms of PV interneuron pathophysiology, relevant to epilepsy and schizophrenia, are distinguished by the level of involvement of PGC-1α.

## Acknowledgments

This work was supported by a Wellcome Trust-NIH PhD studentship (CMGS), and also grants from Epilepsy Research UK (P1504) and Medical Research Council (MR/R005427/1).

## Notes

The authors declare no conflicts of interest

### Competing Interest Statement

The authors have declared no competing interest.

